# Mapping hippocampal-cerebellar functional connectivity across the adult lifespan

**DOI:** 10.1101/2024.10.28.620678

**Authors:** Kavishini Apasamy, Narender Ramnani, Carl J. Hodgetts

**Affiliations:** Department of Psychology, Royal Holloway, University of London, Egham, Surrey, TW20 0EX, UK

**Keywords:** cerebellum, hippocampus, ageing, memory, functional connectivity

## Abstract

Although the hippocampus and cerebellum are traditionally considered to support distinct memory systems, evidence from nonhuman species indicates a close bidirectional relationship during learning and navigational behaviour, with the hippocampus projecting to – and receiving input from – several cerebellar regions. However, little is known about the nature and topography of hippocampal-cerebellar connectivity in the human brain. To address this gap, we applied seed-based functional connectivity analyses to resting-state fMRI data from 479 cognitively normal participants, aged 18–88 years. We identified significant functional correlations between the hippocampus and widespread areas of cerebellar cortex, particularly lobules HIV, HV, HVI, HVIIA (Crus I and II), HIX, and HX. Contrasting the left and right hippocampus, we found significant correlations with the contralateral Crus II. We also compared longitudinal subdivisions of the hippocampus, revealing that anterior hippocampus demonstrated stronger connectivity with right Crus II, whereas posterior hippocampus was strongly connected to vermal parts of lobule V. Finally, we found that functional correlations between several hippocampal seeds (left, right, and anterior) and lobules HVI and HV decreased significantly with age. These results provide novel insights into hippocampal-cerebellar functional organisation and the influence of ageing on this system. Further studies are required to establish the role of this connection in learning and memory, as well as its potential vulnerability to neurodegeneration.

## Introduction

The ability to represent and navigate spatial environments is thought to be supported by an extended posteromedial navigation system in the brain that includes the hippocampus, as well as entorhinal, retrosplenial and parahippocampal cortices (Hodgetts et al., 2017; Murray et al., 2018; Ritchey et al., 2015, see also Ekstrom and Ranganath, 2017). Critically, this brain system is also thought to undergo pronounced structural and functional alterations across the adult lifespan (Lester et al., 2017), acting as a ‘hotspot’ for age-associated neurodegenerative processes, such as tau and amyloid-beta accumulation (Berron et al., 2021; Lace et al., 2009). In this context, it is important to better understand the organisation of the brain’s navigation system, particularly by considering the influence of other cortical and subcortical areas.

The cerebellum, for instance, is classically associated with motor processing (Glickstein, 2007; Glickstein, 1993; Jimsheleishvili and Dididze, 2021) but has been shown to be strongly connected with higher-order cognitive areas, such as prefrontal cortex (Kelly and Strick, 2003; Middleton and Strick, 2001; O’Reilly et al., 2009). In line with this, evidence from nonhuman species suggests that the cerebellum has functional and structural interactions with the hippocampus (Rochefort et al., 2013; Rondi-Reig et al., 2022). For example, electrophysiological studies in transgenic mice have shown that disrupting cerebellar cells alters hippocampal place cell firing and impairs navigational performance (Lefort et al., 2015; Rochefort et al., 2011). Similarly, optogenetic excitation or inhibition of midline and lateral cerebellar neurons has been shown to reduce the duration of left hippocampal seizure activity (Krook-Magnuson et al., 2014), and combined optogenetic stimulation-fMRI revealed that cerebellar stimulation (between lobule V and VI) increased dorsal hippocampal blood-oxygen level dependent (BOLD) signal (Choe et al., 2018). Taken together, these studies establish an argument that the cerebellum modulates the physiology of the hippocampus, as well as influencing hippocampal-dependent behaviour (e.g., spatial learning and memory).

While early electrophysiological evidence in primates and cats suggested a direct structural monosynaptic connection between these structures (as evidenced by short observed latencies between cerebellar fastigial nucleus stimulation and evoked hippocampal responses; Heath and Harper, 1974; Newman and Reza, 1979), recent anatomical tracing work in rodents indicates connectivity through multisynaptic pathways (Watson et al., 2019). Specifically, left hippocampus has been shown to receive input from distributed areas of cerebellar cortex, including bilateral and vermal lobule VI, lobule VIIA (Crus I) and paraflocculus via at least two relay stations (Watson et al., 2019). This study also observed phase-locked theta coherence between cerebellar Purkinje cells in lobule VI, VIIA (Crus I) and the dorsal hippocampus during exploratory behaviour. Notably, evidence in macaque monkeys suggests that lobule VI and VIIA sends its output to the cerebellar fastigial nuclei (Coffman et al., 2011) – the same area found by the above electrophysiological studies to evoke responses in the hippocampus. This suggests that topographically distinct areas of the cerebellum send their input to the hippocampus via distinct pathways.

Despite strong evidence of connectivity in nonhuman species, this is lacking in the human brain. Such evidence has the potential to extend and inform current neurobiological accounts of human memory and navigation, incorporating the cerebellum as a key structure, and may have implications for understanding memory decline in ageing and neurodegeneration.

Studies of healthy ageing in humans, for example, indicate that the cerebellum and hippocampus are both vulnerable to age-related structural changes (Cui et al., 2020; Du et al., 2006; Ramanoël et al., 2023) and exhibit comparable grey matter loss (Bernard and Seidler, 2014; Gellersen et al., 2021; Woodruff et al., 2010). Notably, age-related reductions in cerebellar volume have been linked to poorer performance on hippocampal-dependent navigation tasks (Daugherty and Raz, 2017), suggesting that structural changes may disrupt hippocampal-cerebellar communication. Indeed, recent work in humans has found that age-related cerebellar atrophy is mainly localised to lobule HVI and lobule HVIIA (Crus I; Ramanoël et al., 2023): two regions shown to connect with the hippocampus in nonhuman animal studies (Watson et al., 2019) and co-activate with the hippocampus during navigational learning in young human adults (Iglói et al., 2015). Despite this evidence, there remains a significant gap in understanding hippocampal-cerebellar connectivity in humans, and how it may differ across the lifespan.

Here, we aimed to examine these questions using resting-state fMRI data from the Cambridge Centre for Ageing and Neuroscience (CamCAN) dataset (Taylor et al., 2017). We first sought to map patterns of bilateral hippocampal connectivity (independent of age) within the cerebellum. We hypothesised that the hippocampus would show a strong functional correlation with several areas of the cerebellum, including lobule VI and HVIIA (Crus I), as predicted by prior nonhuman animal tracing work (e.g., Watson et al., 2019) and electrophysiological evidence showing that stimulating fastigial nucleus (which receives input from these areas) leads to evoked responses in the hippocampus (e.g., Coffman et al., 2011). We also contrasted the connectivity of the left hippocampus against the right, drawing on evidence that hippocampally-dependent tasks might be lateralised in the cerebellum (e.g., Riva and Giorgi, 2000). It is also possible that a distinctive pattern of connectivity exists for each hemisphere – a question that remains unresolved as previous anatomical studies have primarily focused on left hippocampal connectivity (e.g., Krook-Magnuson et al., 2014; Watson et al., 2019). The hippocampus is also thought to display functional gradients along its longitudinal (anterior-posterior) axis, arising, in part, from variation in connectional anatomy (Aggleton, 2012; Poppenk et al., 2013; Strange et al., 2014). For instance, the anterior hippocampus preferentially connects to the amygdala and prefrontal cortex (Aggleton et al., 2010; Aggleton, 1986), whereas posterior hippocampus preferentially connects with the parahippocampal cortex (Aggleton, 2012). As regions within these anterior and posterior hippocampal networks might differentially connect with cerebellar cortex (see e.g., prefrontal connections with lobule HVIIA; Ramnani, 2006), we also predicted connectivity differences when contrasting anterior and posterior hippocampal seed regions. Finally, the large age range within the CamCAN dataset (18-87 years old) enabled us to examine how increasing age influences the degree and pattern of hippocampal-cerebellar functional connectivity. We predicted that the strongest age-related decreases in functional connectivity will be observed in lobules HVI and HVIIA, consistent with their vulnerability to age-related atrophy (see e.g., Gellersen et al., 2021; Ramanoël et al., 2023) and suggested involvement in spatial cognition (Iglói et al., 2015).

## Methodology

### Participants

We used structural and functional MRI data from the Cambridge Centre of Ageing and Neuroscience (CamCAN) study (Taylor et al., 2017). This dataset contained 653 participants (323 males, 330 females, 18-87 years old, mean = 54.3, SD=18.6). About fifty males and females were collected from each decile (deciles: 18-27 years, 28-37 years, 38-47 years, 48-57 years, 58-67 years, 68-77 years, 78-87 years; for more information about participants see Shafto et al., 2014). Participants in the dataset were cognitively healthy, assessed via a score above 25 on the mini-mental state examination (Folstein et al.,1975), and did not have any neurological or psychiatric conditions. Written informed consent for the database was obtained in accordance with the Cambridgeshire Research Ethics Committee, and these secondary data analyses were conducted in accordance with Royal Holloway, University of London Ethics Committee processes.

### Neuroimaging data acquisition

All CamCAN MRI data was acquired using a 3T Siemens TIM Trio Scanner at the Medical Research Council (UK) Cognition and Brain Science Unit using a 32-channel head coil. The data used in this study forms part of a larger scanning protocol (see https://camcan-archive.mrc-cbu.cam.ac.uk/dataaccess/ or Taylor et al., 2017 for more detail). High-resolution structural images were obtained using a T1-weighted magnetisation-prepared rapid sequence (MPRAGE; TE= 2.99 ms; TR= 2250ms; TI = 900 ms; voxel size = 1×1×1 mm; field-of-view = 256×240×192 mm; flip angle = 9°). Resting-state fMRI data was acquired using a T2*-weighted gradient echo planar image (EPI) sequence. Rest (resting in the scanner with eyes closed) consisted of 261 volumes and lasted 8 minutes and 40 seconds (32 slices; TE = 30 ms, TR = 1970ms, voxel size: 3×3×4.44 mm; field-of-view: 192×192 mm).

### Data preprocessing and denoising

Initially, functional and structural MRI data from a random sample of approximately 25% of cases were visually inspected for image quality and motion artefacts by KA (with support from CH). Functional MRI images were pre-processed and denoised using the standard pipeline in the CONN toolbox (RRID:SCR_009550; version 22.a; Nieto-Castanon and Whitfield-Gabrieli, 2022) and SPM12 (RRID: SCR_007037; version 12.7771; Penny et al., 2011) running on Matlab 2022b (The Mathworks Inc, Natick, Massachusetts, USA). Functional MRI data were first realigned and unwarped using SPM12 (Andersson et al., 2001). For this, all scans were co-registered to the first volume using a least-squares approach and a 6-parameter rigid body transformation (Friston et al., 1995). These were then resampled using b-spline interpolation to correct for motion and magnetic susceptibly interactions. Temporal misalignment between slices was corrected using the SPM12 slice-timing correction procedure (Henson et al., 1999; Sladky et al., 2011), which involved sinc temporal interpolation to resample each BOLD timeseries slice to a common mid-acquisition time. Potential outlier scans (based on motion and global signal fluctuation) were identified using conservative (95^th^ percentile) outlier parameters in ART (Artifact Detection Tools; Whitfield-Gabrieli et al., 2011). Specifically, volumes with framewise displacement above 0.5 mm, or global BOLD signal changes above 3 SDs, were flagged as outliers (Nieto-Castanon, 2022; Power et al., 2014). A reference mean BOLD image was then computed for each subject by averaging all scans, excluding outliers. Following this, functional and structural images were separately normalised into standard MNI space and segmented into grey matter, white matter and CSF ‘tissue’ types using the SPM12 unified segmentation and normalization algorithm (Ashburner and Friston, 2005; Ashburner, 2007). Functional and structural images were then resampled to 2 mm and 1 mm isotropic voxels, respectively, following a direct normalization procedure (Calhoun et al., 2017; Nieto-Castanon, 2022) with the default IXI-549 tissue probability map template. Finally, functional data was smoothed using a Gaussian kernel of 5 mm full-width half-maximum (FWHM; e.g., Dombrovski et al., 2020).

Following preprocessing, the fMRI data were next denoised using the default pipeline in the CONN toolbox (Nieto-Castanon, 2020). This involved regressing out noise components using an anatomical component-based noise correction procedure (aCompCor), which included noise components from white matter (5 noise components), CSF (5 noise components), motion parameters (3 translational and 3 rotational and their first order derivatives; Friston et al.,1996), outliers volumes derived from scrubbing (38 factors) (Power et al., 2014), effect of rest and its first order derivatives (2 factors; default setting which removes residual trends/instabilities only at the beginning of the timeseries). These were followed by a bandpass frequency filtering of the BOLD timeseries (Hallquist et al., 2013) between 0.01 Hz and 0.09 Hz (e.g., Stefanov et al., 2020), which filters low frequencies (e.g., physiological noise). From the number of noise terms included in this denoising strategy, the effective degrees of freedom of the BOLD signal after denoising were estimated to range from 62.1 to 74.1 (average 70.5) across all subjects (Nieto-Castanon, 2022; see Morfini et al., 2023 for a full description of quality control measures, including degrees of freedom).

Following quality assurance, 479 participants (242 males, 237 females, 18-87 years old, mean= 50.7, SD= 18.2) were entered into the seed-based connectivity analysis. The remaining cases were rejected for reasons such as having more than 10% of invalid scans, motion above 0.5 mm, etc.

### Anatomical methods

Regions of interests (ROIs) for the hippocampus were defined using probabilistic, anatomical atlases. Hippocampal seed ROIs were created by combining the hippocampal ROI from the Harvard-Oxford subcortical atlas and the subiculum ROI from the Jülich histological atlas (Amunts et al., 2005) ensuring that the hippocampal ROI was extended medially incorporating the subicular complex (see Hodgetts et al., 2017). The Harvard-Oxford atlas ROI was thresholded at 50% and the Jülich atlas ROI thresholded at 75% to ensure both ROIs were constrained to grey matter and did not extend into adjacent regions. Using this method, left and right hippocampal ROIs were defined (Figure 1a). For the long-axis analysis, the hippocampal ROIs were split into anterior and posterior zones arbitrarily at the uncal apex (Hodgetts et al., 2017; Poppenk et al., 2013), corresponding to MNI slice y = -21 (see Figure 1b for segmentation of the left hippocampus).

**Figure 1.**
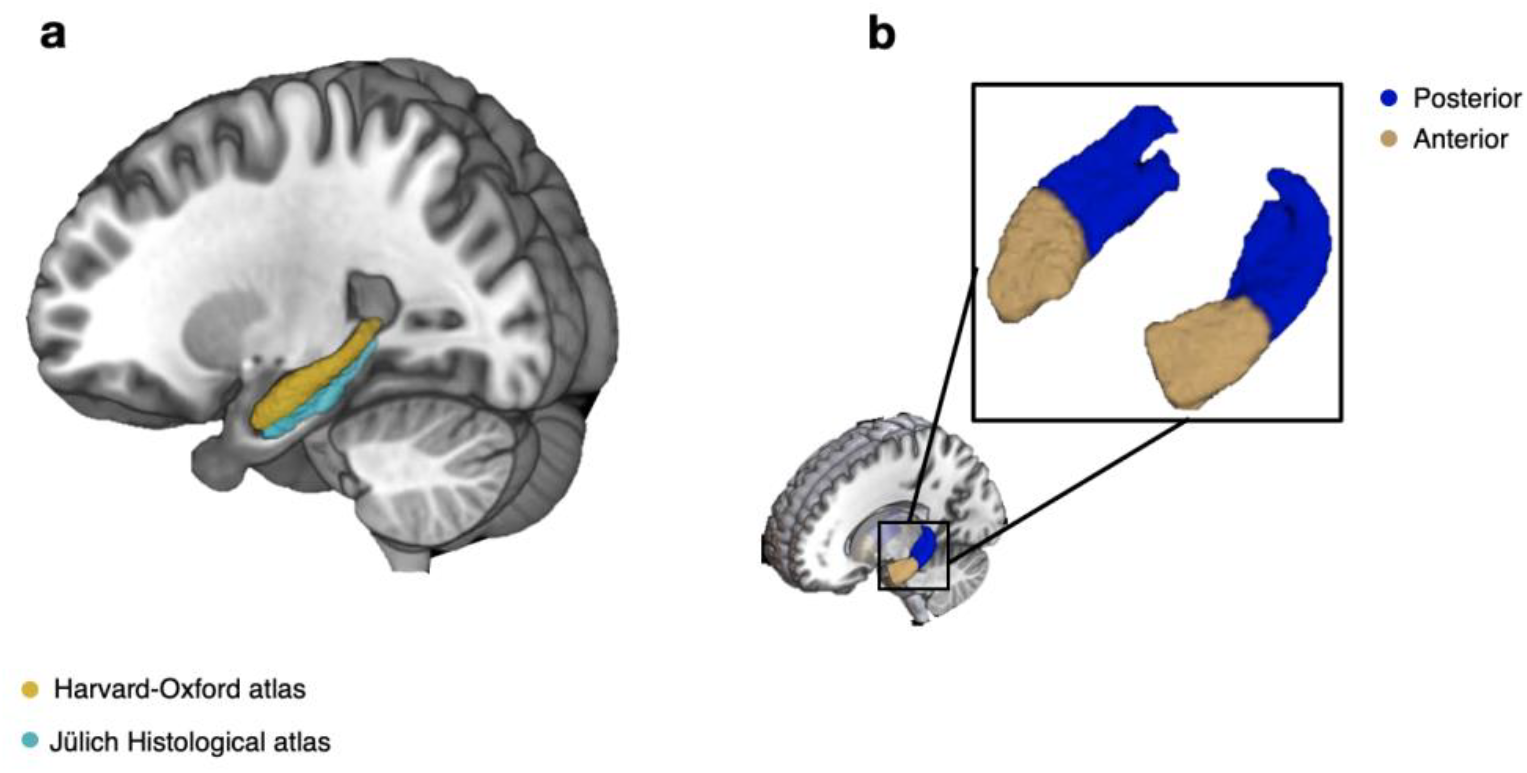
Hippocampal regions-of-interest. (a) The left hippocampal ROI made from combining the hippocampus from the Harvard-Oxford atlas and the subicular complex from the Jüelich Histological atlas; (b) The left and right hippocampal ROI split into anterior and posterior hippocampus using the uncal apex for a landmark-based segmentation.

Functional correlations within the cerebellum were visualised and interrogated using the Spatially Unbiased Infratentorial Template (SUIT) (Diedrichsen et al., 2009). As well as containing a detailed probabilistic atlas of the cerebellum, SUIT also contains a cerebellar flatmap, allowing for visualisation of functional connectivity maps within and across cerebellar lobules/subregions. The nomenclature of Larsell and Jansen (1972) was used to characterise cerebellar cortical results.

### MRI analysis

Following denoising (see above), a first-level seed-based connectivity analysis was conducted using the CONN toolbox. Here, the BOLD timeseries for each hippocampal seed (left hippocampus, right hippocampus, anterior hippocampus, posterior hippocampus), as well as confound regressors (e.g., white matter and CSF) were entered as first-level covariates into general linear models (GLMs). Functional connectivity between the seeds and every other voxel in the brain was represented by the Fisher-transformed bivariate correlation coefficient (*r*-to-Z) from a weighted GLM. At the first level, we conducted eight analyses whereby four of them related to single seed and four of them to between-seed contrasts. Here we considered the issue of shared variance, and this was minimised by the use of separate GLMs for each seed which avoids multicollinearity between seeds in the same model. For the single seed analyses, the left, right, anterior, and posterior hippocampus design matrices contained a single column representing all subjects, allowing the investigation of seed-to-voxel connectivity for each ROI in isolation.

To examine hemispheric and long-axis differences in hippocampal connectivity with the cerebellum, we specified between-seed contrasts (referred to as between-source contrasts in CONN). Here, the design matrices included two columns representing each seed in each contrast. Contrasts compared the connectivity of left and right hippocampus (contrast vectors: 1 -1 and -1 1). Others compared the connectivity of anterior and posterior hippocampus (contrast vector: 1 -1 and -1 1). For single seed analyses, beta images were carried to the second-level analysis for one-sample *t*-tests. For between-seed analyses, contrast images generated in the first-level analysis were carried over to the second-level analysis for one-sample *t*-test. To explore age-related variation in hippocampal-cerebellar functional correlations, we also re-specified these GLMs with age as an additional subject-level effect, firstly as a regressor-of-no-interest (contrast vector: 1 0), and then as a regressor-of-interest (contrast vector: 0 1) into a bivariate regression analysis. For these contrasts, age was mean-centred and to ensure that we captured anticorrelations, we applied a contrast vector of 0 -1 to the demeaned age data.

The unthresholded group-level *t*-statistic maps (as well as between-source contrast maps) from CONN were interrogated in SPM. These were thresholded using a family-wise error correction *p-*FWE *<*0.05 based on Random Field Theory (Worsley et al., 1996). The thresholded images were then loaded into SUIT to localise (and visualise) significant clusters within cerebellar subregions. This produced cerebellar flatmaps showing suprathreshold connectivity strength within and across cerebellar lobules.

## Results

### The hippocampus is functionally connected to widespread areas of the cerebellar cortex

Initially, we used seed-based connectivity to examine patterns of cerebellar connectivity from left and right hippocampal seed ROIs. As predicted by anatomical studies, both the left and right hippocampus showed bilateral connections to widespread areas of the cerebellum, including the vermis and lateral hemispheres. Both hemispheres showed strongest connections with the border of lobule HIV and HV, HIX, HX, and lobule HVIIA (medial parts of Crus II and laterally within the horizontal fissure at the junction of Crus I and Crus II; Figure 2a-b).

**Figure 2.**
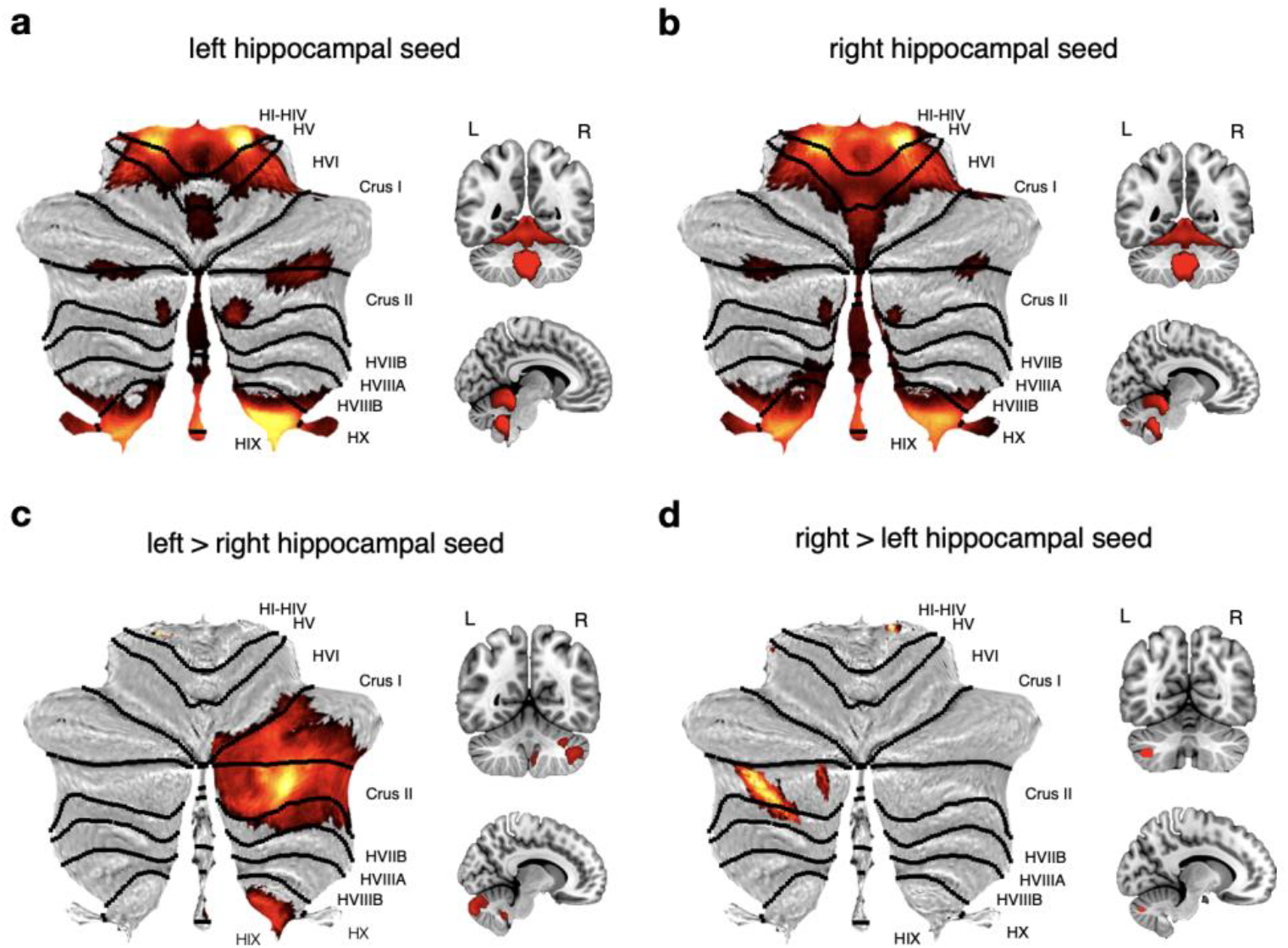
Cerebellar regions showing significant functional connectivity with left and right hippocampus. Thresholded SPM {T} maps (*p*<0.05) overlaid on the cerebellar flatmap using SUIT (Diedrichsen et al., 2006) and on coronal and mid-sagittal sections of the standard MNI T1 2mm template. Contrasts for main effects of LEFT and RIGHT hippocampal seeds (a) and (b), respectively; contrasts for LEFT > RIGHT and RIGHT > LEFT hippocampal seeds, (c) and (d), respectively.

To examine hemispheric differences, cerebellar functional correlations with left and right hippocampus were contrasted directly. The left hippocampus, compared to the right hippocampus, showed preferential connections with a contralateral region of lobule HVIIA (Crus I and peak in Crus II; number of voxels = 3063; *p*<.001), as well as lobule HIX (number of voxels = 181, *p*<.001; Figure 2c). Similarly, the right hippocampal seed also showed a strong preferential functional correlation with the contralateral (i.e., left) region of lobule HVIIA (peak in Crus II; number of voxels = 178, *p*<.001; Figure 2d). Peak cluster statistics for these hemispheric contrasts are shown in Table 1.

**Table 1.**
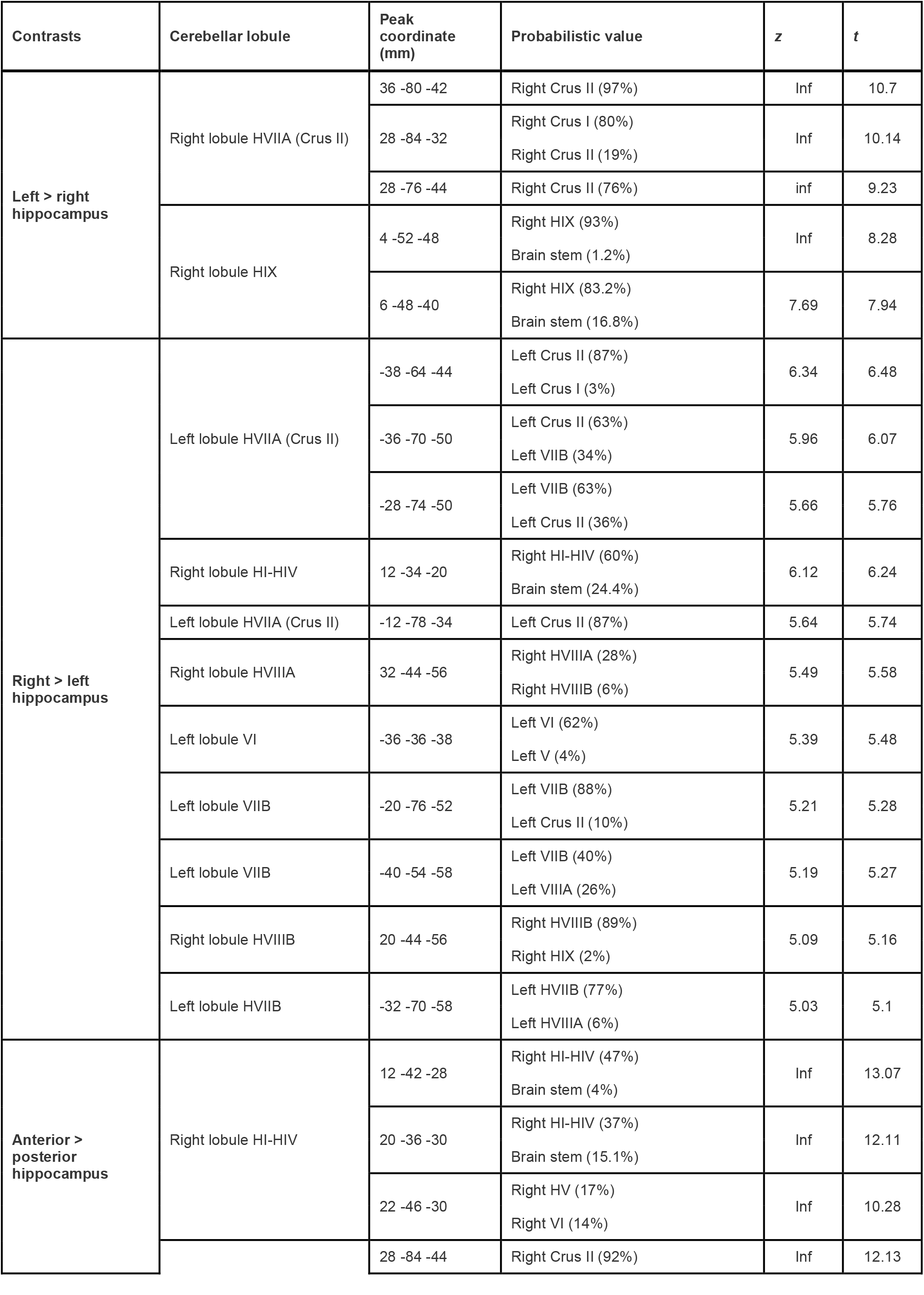
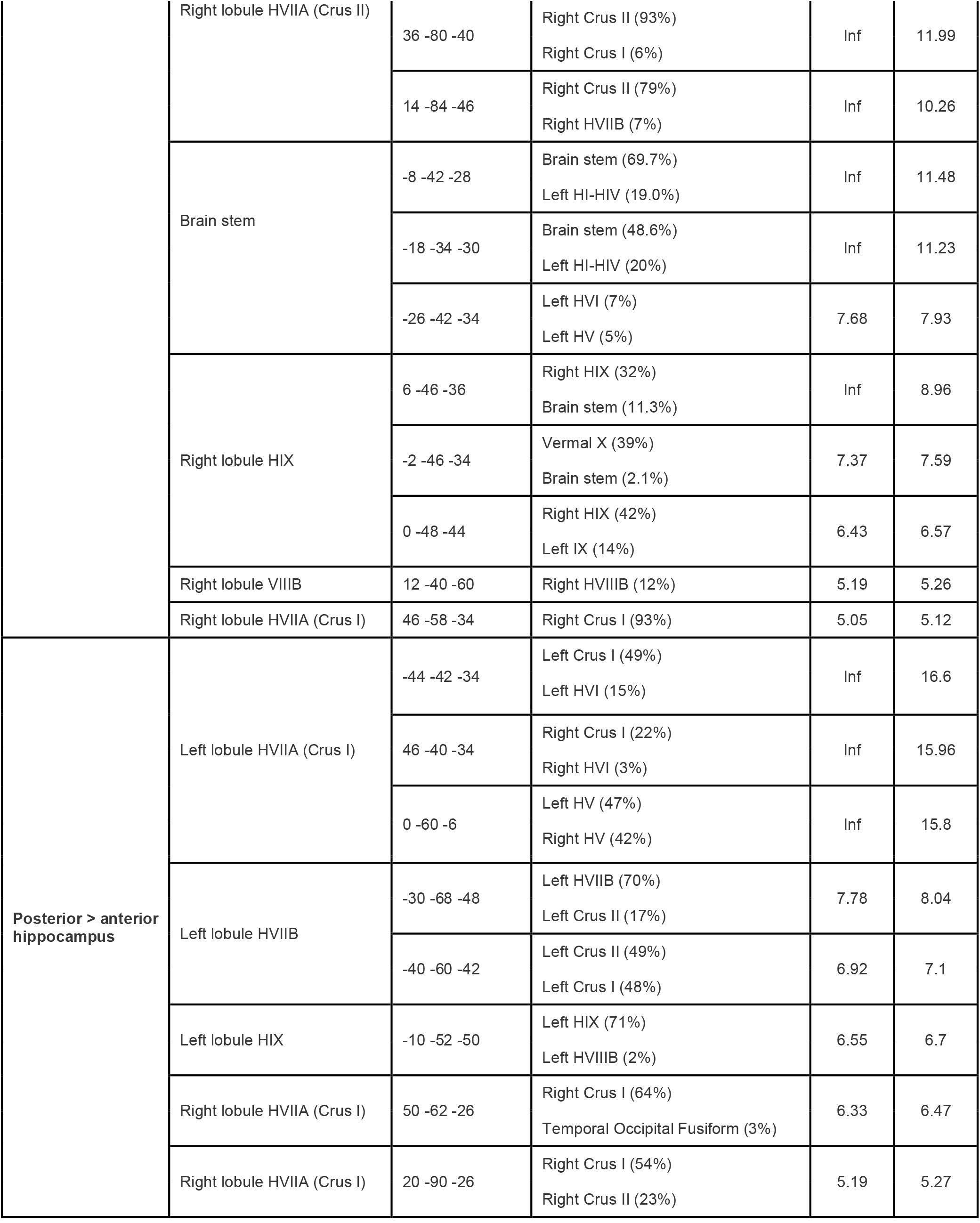
Significant clusters identified within the cerebellum. Peak coordinates, z- and t-values are reported for all clusters. All statistics are reported at FWE *p*<0.05. NOTE: Coordinates are reported in MNI 2 mm space. Negative X-coordinates indicate the left hemisphere. Inf indicates large values that surpass threshold. Probabilistic values are derived from SPM Anatomy Toolbox (Eickhoff et al., 2005).

### Long-axis subdivisions of the hippocampus show both overlapping and distinct patterns of cerebellar connectivity

Next, we examined hippocampal-cerebellar functional connectivity along the longitudinal axis of the hippocampus. The anterior and posterior hippocampus showed connectivity to similar regions as those connected to the left and right hippocampus, including the border of lobule HIV and HV, vermal parts of lobule V, lobule HVIIA (bilaterally at the fissure separating Crus I and II), lobule HIX and HX (Figure 3a-b).

**Figure 3.**
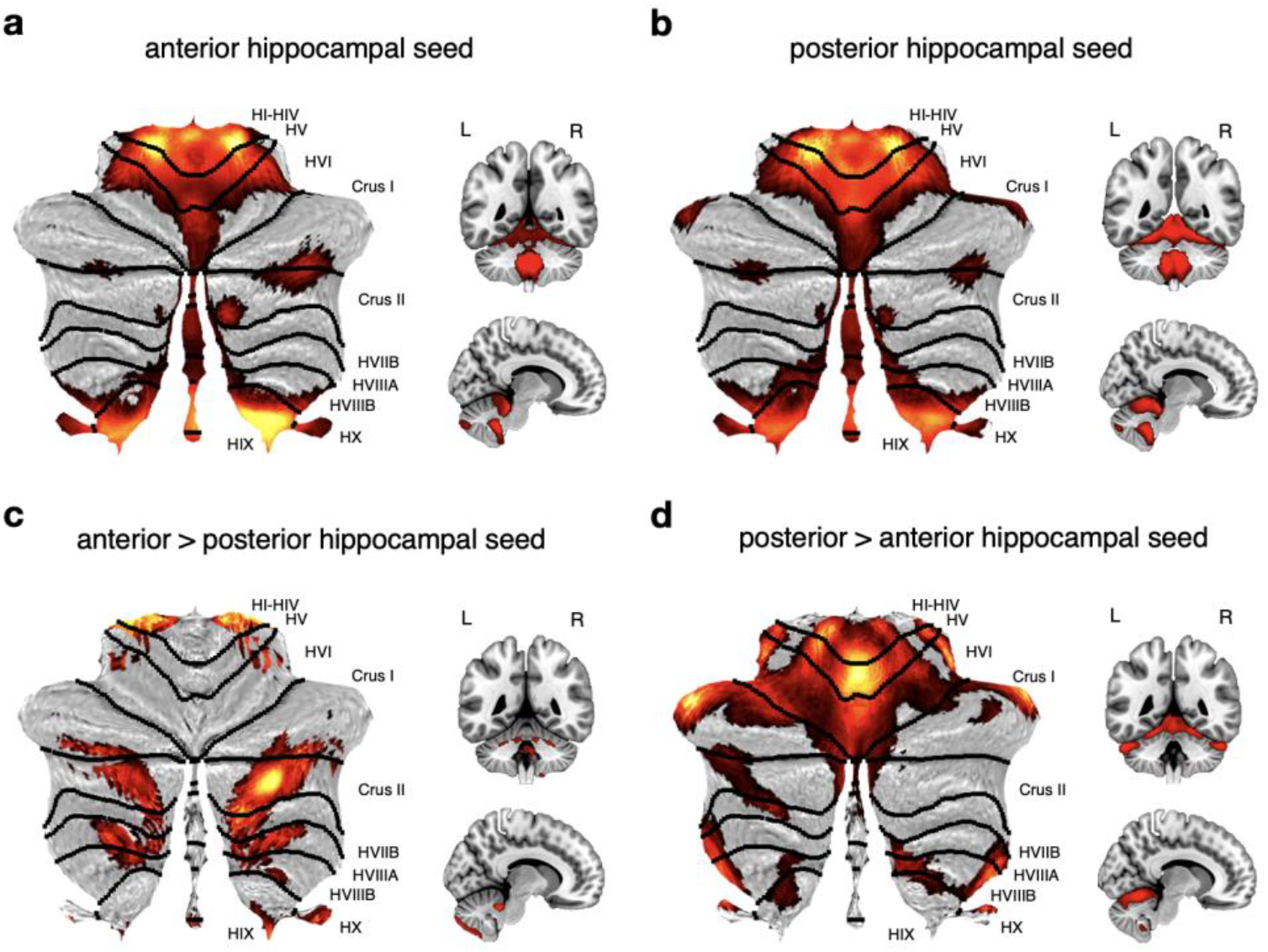
Cerebellar regions showing significant functional connectivity with anterior and posterior hippocampal seeds. Thresholded SPM {T} maps (*p*<0.05) overlaid on coronal, sagittal slices and cerebellar flatmaps. Contrasts for main effects of ANTERIOR and POSTERIOR hippocampal seeds (a) and (b), respectively (c) represents contrast for ANTERIOR > POSTERIOR hippocampal seed connectivity with the cerebellum. (d) represents contrast for POSTERIOR > ANTERIOR hippocampal seed connectivity with the cerebellum.

However, direct contrasts revealed that the anterior hippocampus, compared to the posterior hippocampus, showed significantly greater functional correlations with lobule HI-HIV (number of voxels = 185, *p*<.001) and bilateral regions of lobule HVIIA, including right Crus II (number of voxels = 2778, *p*<.001; Figure 3c). Note, this same region of Crus II also displays significantly greater connectivity with left versus right hippocampus (Figure 2c). Additional clusters were also seen in extreme areas of right lobule HIX and HX, and bilateral HVIIB and HVIII. The posterior versus anterior hippocampus contrast showed significantly greater connectivity with widespread regions of the anterior cerebellum. Specifically, we observed a peak in the bilateral extremes of the Crus I region of lobule HVIIA (number of voxels = 4761; *p*<.001), which extended medially into vermal areas of lobule V (see Figure 3d). Further small clusters were found in the lateral extremes of lobule HVIIIA, bilaterally, which continued along the border of HVIIIB and HIX. Peak cluster statistics for these direct long-axis contrasts are shown in Table 1.

### Hippocampal-cerebellar functional connectivity is reduced in ageing

Finally, we examined the effect of age on hippocampal-cerebellar functional connectivity. We observed that several cerebellar regions showed negative correlations between age and hippocampal-cerebellar functional connectivity (i.e., lower connectivity in older participants). For the left hippocampal seed, we saw large age-related reductions in the depths of the right primary fissure that separates cerebellar lobules HV and HVI (number of voxels = 382; *p*<.001), which extended into adjacent parts of lobule HVIIA (Crus I; see Figure 4a). A similar spatial pattern of connectivity alterations was seen for the right hippocampus, but also revealed a strong cluster in the contralateral region of lobule HVI (number of voxels = 71; *p*<.001 (see Figure 4b). No differences were observed when directly contrasting left and right hippocampus.

**Figure 4.**
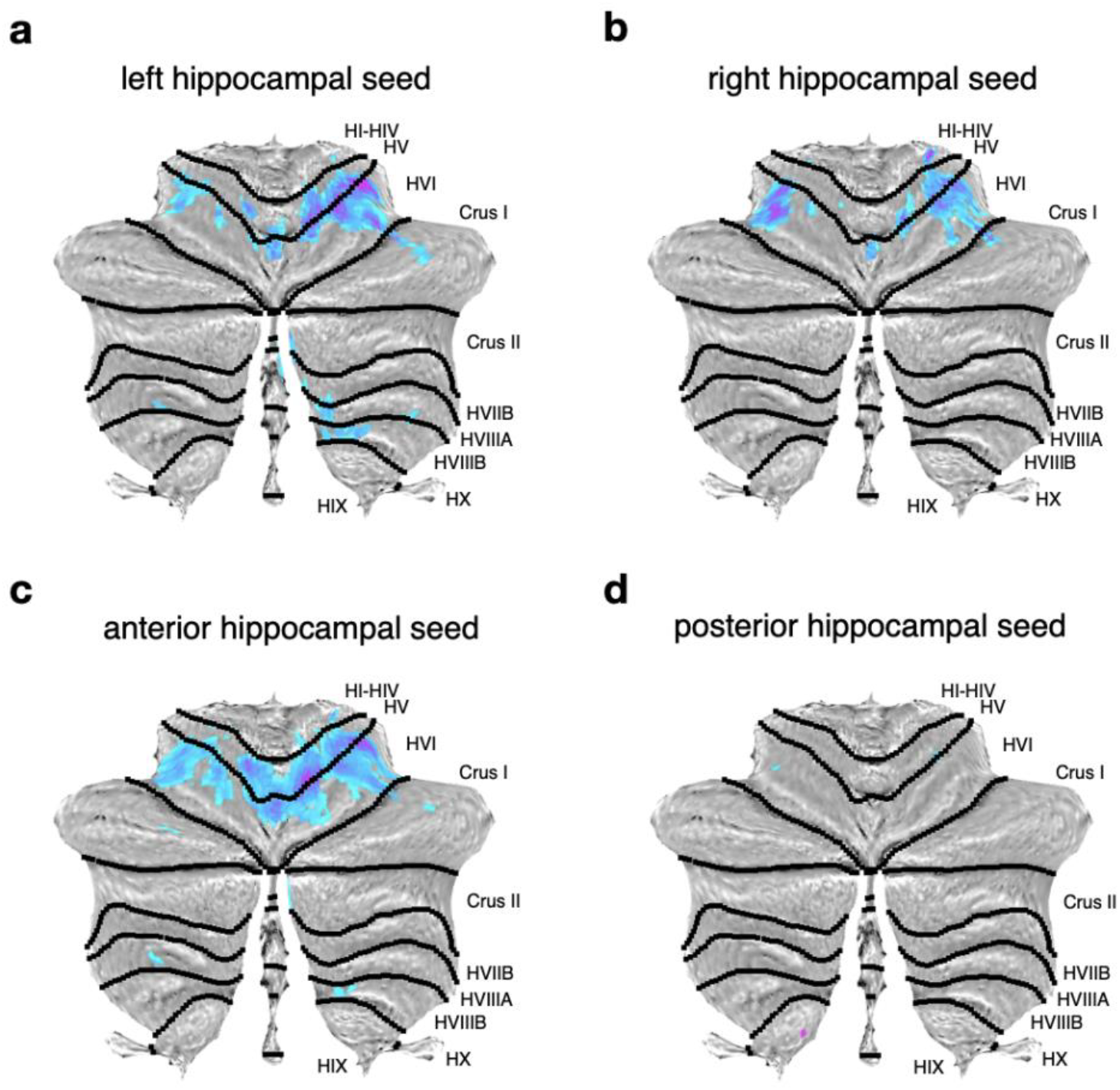
Regions in cerebellar cortex showing age-related decreases in functional connectivity with key hippocampal seeds. (a) Thresholded SPM {T} maps (*p*<0.05) overlaid on cerebellar flatmaps showing ageing main effect for left (a), right (b), anterior (c) and posterior (d) hippocampus to the cerebellum. Darker colours indicate *decreased* functional connectivity with seed ROI with age.

Next, we explored whether these age-related effects within the cerebellum differed across different hippocampal long-axis subdivisions. Notably, we found that the anterior hippocampus showed widespread age-related connectivity reductions within the cerebellum, and these were present primarily within the primary fissure between lobules HV and HVI (number of voxels = 837; *p*<.001; see Figure 4c). In contrast, the posterior hippocampus showed minimal age-related changes in functional connectivity, with very small suprathreshold clusters observed in dispersed areas of the cerebellum, including lobules HV and HVI, with a peak in lobule HIX (number of voxels = 1; *p* = .026; see Figure 4d). However, no age-dependent connectivity differences were observed when contrasting anterior and posterior hippocampus. Cluster statistics for the ageing analyses are shown in Table 2.

**Table 2.**
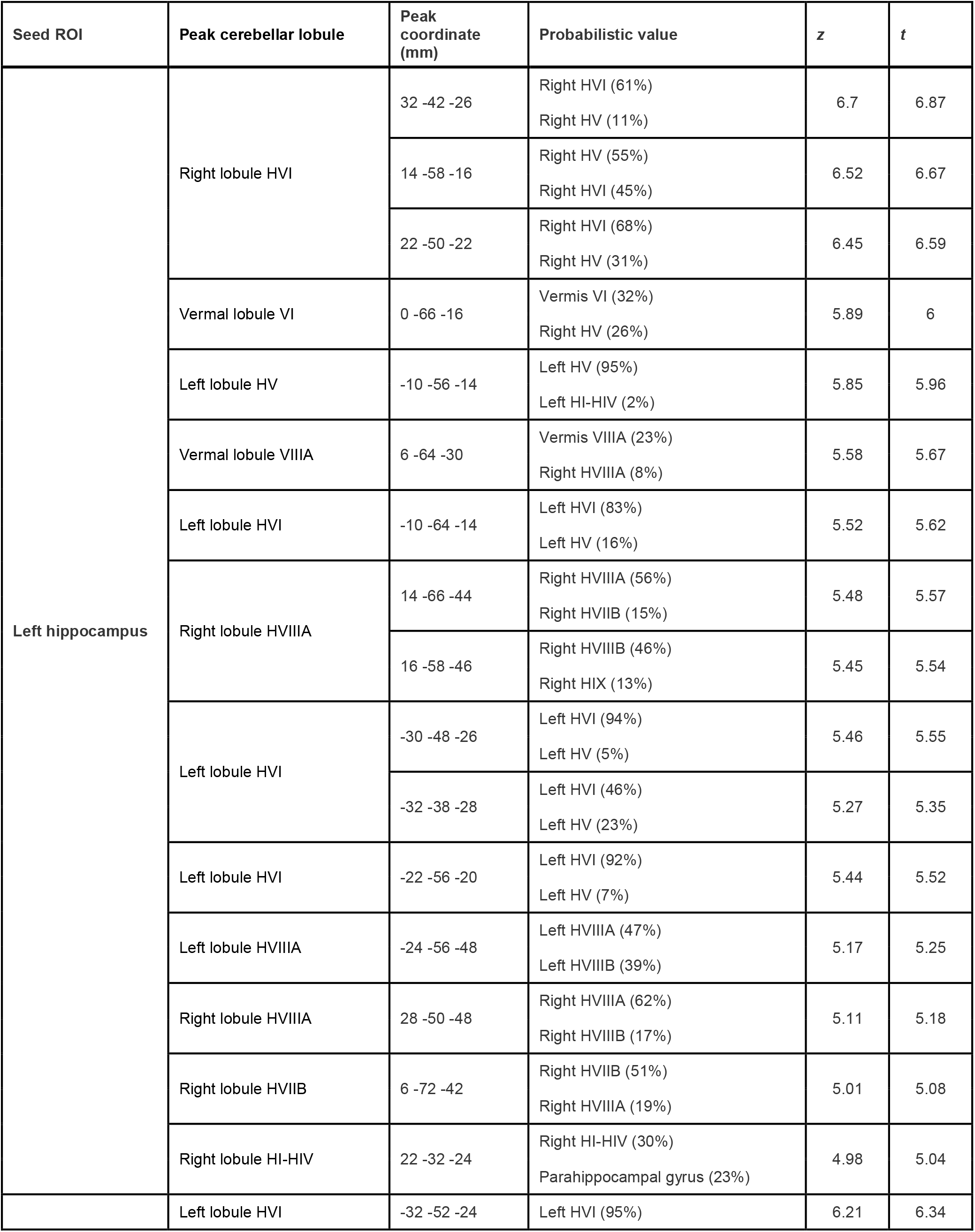
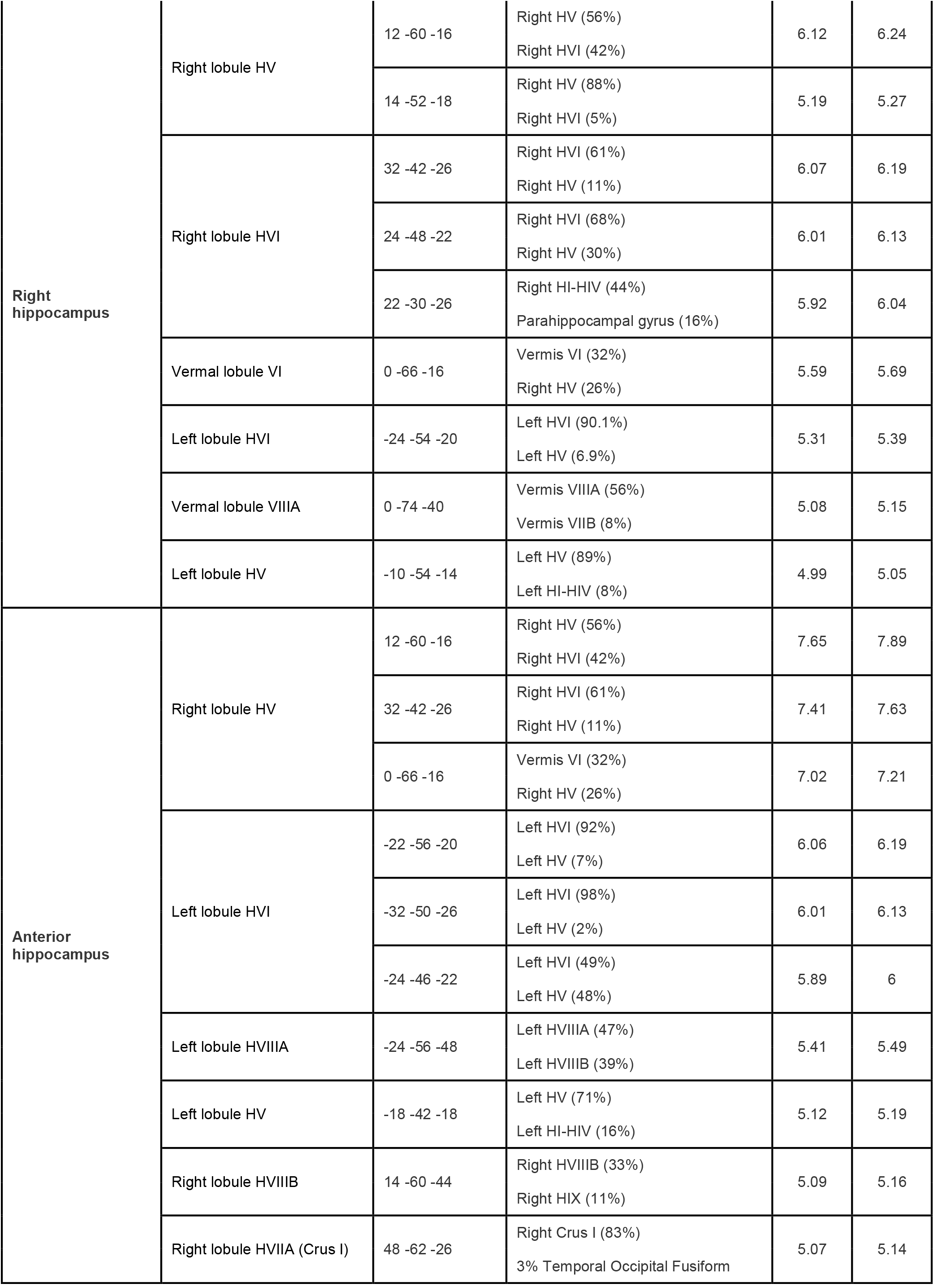
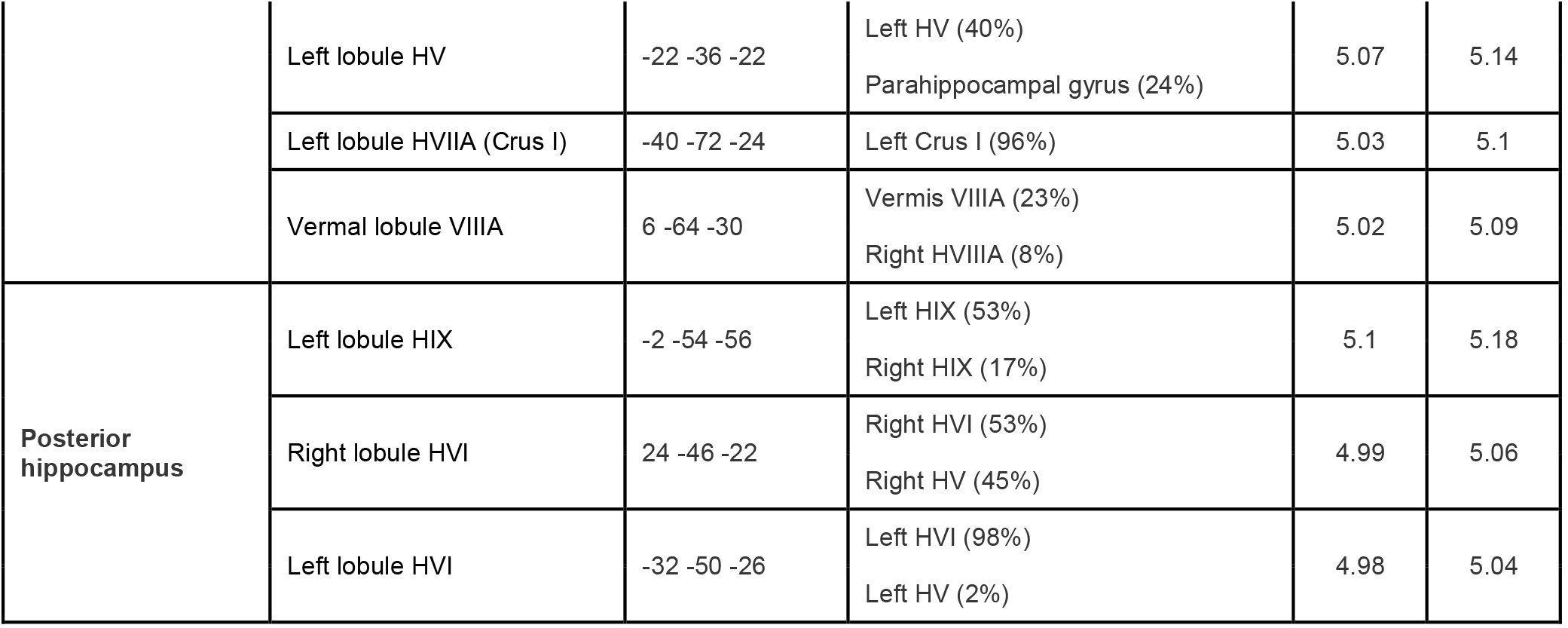
Significant clusters displaying significant age-related reductions in functional connectivity in the cerebellum. Peak coordinates, z- and t-values are shown for all clusters. All statistics are reported at FWE *p*<0.05. NOTE: Coordinates are reported in MNI 2 mm space. Negative X-coordinates indicate the left hemisphere. Probabilistic values are derived from SPM Anatomy Toolbox (Eickhoff et al., 2005).

## Discussion

While prior evidence in nonhuman animals suggest that a close functional interaction exists between the hippocampus and cerebellum, there is a limited understanding of the nature and topography of this functional connection in humans and how it be might vary with age. In the current study, we sought to close this gap by applying seed-based functional connectivity analyses to resting-state fMRI data collected in a large-scale lifespan cohort (N = 479 from CamCAN). Our results yielded several key findings. Firstly, the shared bilateral functional connectivity between the left and right hippocampus with cerebellum involves four areas of the cerebellar cortex bilaterally: (i) almost the whole of the anterior lobe, and in addition, the anterior bank of lobule HVI; (ii) lobule HVIIA, both the anterior and posterior bank in the mid-point of the horizontal fissure; (iii) a separate region in lobule HVIIA in Crus II, adjacent to a medial part of the ansoparamedian fissure; and (iv) lobules HIX and HX. We also considered laterality-related connectivity differences. When directly contrasting the left and right hippocampus, we found that the left hippocampus showed greater connectivity with contralateral lobules HVIIA (Crus I and Crus II) and HIX, whilst the right hippocampus showed greater connectivity with the contralateral lobule HVIIA (Crus II).

Second, we found that the shared functional connectivity of the anterior and posterior hippocampus with the cerebellum encompassed similar areas as the bilateral shared functional connectivity of the left and right hippocampus. When examining differential functional connectivity between anterior and posterior hippocampus, we found that these individually correlated with distinct parts of the cerebellar cortex. The anterior compared to posterior hippocampus contrast revealed largely medial cerebellar connectivity dominated by areas posterior to the horizontal fissure (including the mid-portion of lobule HVIIA, extending into more medial parts of HVIIB and HVIII and small areas of lobule HIX and HX). The posterior compared to anterior hippocampus contrast revealed connectivity with areas of the cerebellar cortex that were largely anterior to the horizontal fissure, including the vermal portion of lobules I to VII, and hemispherical portions of lobules HI to HVIIA, including Crus I.

Finally, we observed age-related decreases in hippocampal-cerebellar connectivity that were confined largely to areas of the anterior lobe. Similar patterns were observed for left and right hippocampus, whereby decreases were observed in lobule HVI, and around the right primary fissure. Anterior hippocampus showed age-related decreases in connectivity with those same areas of the cerebellar cortex, whereas the posterior hippocampus revealed no appreciable effects in any part of it.

Previous electrophysiological studies have suggested a strong functional interaction exists between the hippocampus and cerebellum, based on the effects of cerebellar disruption on hippocampal place cell firing (Lefort et al., 2015; Rochefort et al., 2011) and hippocampal-cerebellar synchronous oscillatory activity (Watson et al., 2019). The current findings support notions that these functional interactions exist in the human brain but, importantly, elucidates the topography/spatial organisation of these functional interactions in the cerebellum. The findings of hippocampal-cerebellar connectivity within cerebellar lobule HIV, HV, HVI and bilateral regions of lobule HVIIA (Crus I and Crus II) aligns with prior electrophysiological findings (e.g., Babayan et al., 2017; Watson et al., 2019; Zeidler et al., 2020) and anatomical work in mice (e.g., Bohne et al., 2019; Watson et al., 2019). For instance, hippocampal field potentials have been shown to be modulated, most strongly, by stimulation of (vermal) lobules IV-V (Zeidler et al.,2020), and phase-locked theta coherence has been observed between dorsal hippocampus and cerebellar lobule VI and VIIA (Crus I; Watson et al., 2019). Likewise, the combination of c-Fos imaging and graph theory suggest that two cerebellar networks – one including lobule IV-V, VI, lobule VIIA (Crus I), and other including lobule IX, X – are differentially connected with dorsal hippocampal CA1 neurons during spatial exploration (Babayan et al., 2017). Notably, these same cerebellar lobules are also functionally correlated with left, right, anterior and posterior hippocampal seeds in this study (see also Figure 2a-b, 3a-3b).

Our findings also align with previous tracing studies. For example, tracing injections into the dentate gyrus of the mouse hippocampus identified polysynaptic connectivity with lobule HIV and lobule HV (Bohne et al., 2019). Further, retrograde tracing studies also identified inputs to the left hippocampus from lobule HVI and lobule HVIIA (Crus I) (Watson et al., 2019) – all regions which were also found be functionally connected with the hippocampus in the current study, some of these even showing preferential connections with specific hippocampal regions (e.g., lobule HIV, HV and HVI with posterior hippocampus).

Our findings also provide novel insights into hippocampal-cerebellar connectivity in humans. For example, in contrast to prior work in nonhuman species (discussed above), the strongest connectivity in lobule HVIIA was found in Crus II rather than the more commonly reported Crus I (e.g., Watson et al., 2019). This was particularly evident when contrasting anterior and posterior hippocampus (a comparison which has not been made in the animal studies). Specifically, while each long-axis subdivision showed similar patterns of cerebellar connectivity when considered in isolation (and indeed showed patterns akin to the hippocampus as a whole), when directly contrasted we found that the anterior hippocampus was strongly correlated with Crus II (extending into lobule HVIIB and HVIII). In comparison, the posterior hippocampus showed the strong functional correlation with lobule HVIIA and extending HV (extending into lobule VI). In fMRI studies, Crus II has been shown to be involved in more abstract, non-motor functions, such as first- and second-order rule learning (Balsters et al., 2013), creative thinking (Gao et al., 2020), and mentalising (Guell and Schmahmann, 2020; Van Overwalle et al., 2020). In contrast, the regions identified in the posterior versus anterior contrast (notably lobule V) are implicated in sensorimotor processing (e.g., Bushara et al., 2001) and fine-grained digit representations (Grodd et al., 2001; van der Zwaag et al., 2013). The differential connectivity of anterior and posterior hippocampus to functionally distinct regions of the cerebellum is interesting as it suggests that functional subdivisions within the hippocampal formation – which are considered to emerge partially from differences in neocortical connectivity (Adnan et al., 2015; Aggleton, 2012; Dalton et al., 2019) – are also reflected within the cerebellum. The cerebellum is thought to have motor and non-motor loops with different cortical areas (Ramnani et al., 2006). Given our data, it is possible that the anterior hippocampus participates more strongly in the cerebellar non-motor loop (consistent with its increased connectivity with prefrontal cortex), whilst the posterior hippocampus participates more strongly in the cerebellar motor loop. Such a distinction aligns with current views of long-axis specialisation in the hippocampus, in which anterior hippocampus provides more abstract or coarse-grained representations in spatial and episodic memory, whereas the posterior hippocampus supports more fine-grained spatial processing (Poppenk et al., 2013; Robin and Moscovitch, 2017).

Our findings also suggest that hippocampal-cerebellar connectivity decreases with age, and this was most evident in areas of cerebellar lobules HV, HVI and their border. This finding dovetails with previous studies that have observed structural and functional changes in both the hippocampus and the cerebellum throughout ageing, making it plausible that this reflects, in part, age-related alterations in hippocampal-cerebellar interactions. Indeed, the current findings show that the left, right and anterior hippocampus show age-related connectivity reductions with areas of cerebellar lobules HV, HVI and their border. This evidence is in accordance with previous evidence that reported atrophy in lobule HVI during ageing (Cui et al., 2020; Ramanoël et al., 2023). Reduced grey matter volume in lobule HVI was also negatively associated with performance on a perspective-taking task (Ramanoël et al., 2023) – a task which is thought to involve hippocampally-dependent allocentric representations (Labash et al., 2020). Further, fMRI work in young adults reported increased but differential activation for place-versus sequence-based navigation in hemispheric lobule HVI (Iglói et al., 2015). The age-related reduction in hippocampal-lobule HVI connectivity seen here, therefore, may partly reflect age-related changes in the ability to adopt allocentric or place-based strategies during navigation, as observed in other studies (Moffat and Resnick, 2002; Rodgers et al., 2012).

In this context, it is important to note that while our results strongly support a close functional interaction between hippocampus and cerebellum, they cannot speak directly to how such connectivity supports spatial/mnemonic behaviour. In nonhuman species, there is growing evidence that the cerebellum directly influences hippocampal spatial representations and navigational behaviour (e.g., Rochefort et al., 2013). One particular fMRI investigation in humans found that distinct hippocampal-cerebellar connectivity patterns might relate to the application of different strategies during virtual navigation. For instance, it was found that the right hippocampus and contralateral lobule HVIIA (Crus I) co-activated during place-based navigation whilst left hippocampus and contralateral lobule HVIIA (Crus I) co-activated to support sequence-based navigation (Iglói et al., 2015). These same cerebellar regions were also strongly functionally correlated with the hippocampus in this study.

One possible account for these interactions is that the cerebellum supports the updating and re-organisation of hippocampally-based spatial representations (e.g., cognitive maps) via its use of forward models (Ramnani, 2014; Rondi-Reig et al., 2022). This describes a model, created from an internal command, that is an internal representation of the behaviour modified by sensory feedback (Ito, 2008). Upon receiving sensory information that mismatches the internal representation, the cerebellum could support the re-computation of cognitive maps in the hippocampus when novel information is encountered.

An additional finding in the current study was that several hippocampal seeds showed strong functional correlations with regions of the ‘vestibulocerebellum’ – namely, lobules IX and X. While speculative, this functional interaction is consistent with prior work in rodents that has demonstrated the importance of vestibular information for the accuracy and stability of hippocampal representations (Sharp et al., 1995), and aligns with theoretical models that suggest that the cerebellum supplies the hippocampus with information in an appropriate (world-centred) format to be used for allocentric navigation (Rochefort et al., 2013).

Functional connectivity approaches also cannot elucidate how the hippocampus and cerebellum are *structurally* connected. Whilst it was previously speculated that a direct route exists between these regions (Heath and Harper, 1974), recent work points towards a more indirect route via a number of relay stations. These include thalamic regions (e.g., laterodorsal and ventrolateral thalamus; Bohne et al., 2019), which receive input from the cerebellum (Fujita et al., 2020) and project to areas that input to the hippocampus, such as the retrosplenial cortex (Van der Werf et al., 2002). Other relay stations include the septum and supramamillary nucleus, which are thought to be key regions involved in generating theta, and thus potentially relevant to theta coupling observed between hippocampus and cerebellum during navigation/exploration (Watson et al., 2019) and cerebellar-dependent eye-blink conditioning (Hoffmann and Berry, 2009). It may be possible to map structural pathways between cerebellar regions/nuclei and the human hippocampus by leveraging high-resolution diffusion MRI tractography, though the range and complexity of fiber populations (e.g., through the midbrain) would require advanced modelling approaches and potential validation data in other species. One study, for example, used probabilistic constrained spherical deconvolution tractography (which enables complex/crossing fiber populations to be better resolved) and identified tractography streamlines between the hippocampus and cerebellar lobules HVIII, IX, X, Crus I, Crus II and the fastigial nucleus (Arrigo et al., 2014).

Here, through the application of seed-based functional connectivity analyses within a large lifespan imaging cohort, we demonstrated that the human hippocampus is strongly functionally correlated with widespread areas of the cerebellar cortex, including lobule I-IV, V, VI, Crus I, Crus II, IX and X. This finding provides additional important support for the idea that the two key brain structures, which are classically considered to underpin distinct memory systems, may collaborate closely in the human brain. Further, we show that hippocampal-cerebellar connectivity decreases significantly across the lifespan, particular in cerebellar regions that are vulnerable to age-related structural atrophy (lobules HV and HVI). It will be important for future studies to examine this interaction during behaviour, which will inform the development of more detailed neurobiological models of (spatial) learning and memory that incorporate these distinct regions, as well as examine the vulnerability of this functional link in neurodegenerative disorders (e.g., Alzheimer’s disease).

## Acknowledgements

This work was supported a Sarah Parker Remond PhD studentship at Royal Holloway, University of London, and the Biotechnology and Biological Sciences Research Council (BBSRC) [BB/V010549/1]. For the purpose of open access, the authors have applied a Creative Commons Attribution (CC BY) licence to any Author Accepted Manuscript version arising. We would also like to thank Samuel Berry, Marie-Lucie Read, Eleanor Alderman and Kieran Allen for their valuable advice during this project.

## CRediT author statement

**Kavishini Apasamy:** Conceptualization, Formal Analysis, Data Curation, Visualization, Writing – Original Draft, Writing – Reviewing and Editing. **Narender Ramnani:** Conceptualization, Data Curation, Visualization, Supervision, Writing – Reviewing and Editing. **Carl J. Hodgetts:** Conceptualization, Visualization, Supervision, Writing – Reviewing and Editing, Funding acquisition.

